# PREDICTIVE IMMUNE CHECKPOINT BLOCKADE CLASSIFIERS DISTINGUISHING MONO- VERSUS COMBINATION THERAPY REQUIREMENT

**DOI:** 10.1101/2020.07.14.202408

**Authors:** Oscar Krijgsman, Kristel Kemper, Julia Boshuizen, David W. Vredevoogd, Elisa A. Rozeman, Sofia Ibanez Molero, Beaunelle de Bruijn, Paulien Cornelissen-Steijger, Aida Shahrabi, Martin Del Castillo Velasco-Herrera, Ji-Ying Song, Maarten A. Ligtenberg, Roelof J.C. Kluin, Thomas Kuilman, Petra Ross-MacDonald, John Haanen, David J. Adams, Christian Blank, Daniel S. Peeper

## Abstract

Although high clinical response rates are seen for immune checkpoint blockade (ICB) treatment of metastatic melanomas, both intrinsic and acquired ICB resistance remain considerable clinical challenges^1^. Combination ICB (anti-PD-1 + anti-CTLA-4) shows improved patient benefit^2–5^, but is associated with severe adverse events and exceedingly high cost. Therefore, there is a dire need to stratify individual patients for their likelihood of responding to either anti-PD-1 or anti-CTLA-4 monotherapy, or the combination. Since it is conceivable that ICB responses are influenced by both tumor cell-intrinsic and -extrinsic factors^6–9^, we hypothesized that a predictive genetic classifier ought to mirror both these features. In a panel of patient-derived melanoma xenografts^10^ (PDX), we noted that cells derived from the human tumor microenvironment (TME) that were co-grafted with the tumor cells were naturally replaced by murine cells after the first passage. Taking advantage of the XenofilteR^11^ algorithm we recently developed to deconvolute human from murine RNA sequence reads from PDX^10^, we obtained curated human melanoma tumor cell RNA reads. These expression signals were computationally subtracted from the total RNA profiles in bulk (tumor cell + TME) melanomas from patients. We thus derived one genetic signature that is purely tumor cell-intrinsic (“InTumor”), and one that comprises tumor cell-extrinsic RNA profiles (“ExTumor”). Here we report that the InTumor signature predicts patient response to anti-PD-1, but not anti-CTLA-4 treatment. This was validated in melanoma PDX and cell lines, which confirmed that InTumor^LO^ tumors were effectively eliminated by adoptive cell transfer of T-Cell Receptor (TCR)-matched cytotoxic T cells, whereas InTumor^HI^ melanomas were refractory and grew out as fast as tumors challenged with unmatched T cells. In contrast, the ExTumor signature predicts patient response to anti-CTLA-4 but not anti-PD-1. Most importantly, we used the InTumor and ExTumor signatures in conjunction to generate an ICB response quadrant, which predicts clinical benefit for five independent melanoma patient cohorts treated with either mono- or combination ICB. Specifically, these signatures enable identification of patients who have a much higher chance of responding to the combination treatment than to either monotherapy (p < 0.05), as well as patients who are likely to experience little benefit from receiving anti-CTLA-4 on top of anti-PD-1 (p < 0.05). These signatures may be clinically exploited to distinguish patients who need combined PD-1 + CTLA-4 blockade from those who are likely to benefit from either anti-CTLA-4 or anti-PD-1 monotherapy.

## RNA signals from tumor cells and TME can be computationally dissected

Anti-CTLA-4 and anti-PD-1 treatments differentially affect tumor and TME^6,7^, and hence we argued that both effects should be accounted for when building gene expression signatures to predict clinical response. To distinguish between tumor cell-intrinsic and -extrinsic gene expression, we exploited our previously established stage IV metastatic melanoma PDX cohort (n=95)^10^. Computational analysis comparing melanoma RNA expression of both tumor and TME genes from patients (from TCGA, n=367) with genes from these PDX demonstrated that mouse TME had almost completely replaced human TME upon PDX outgrowth after the first passage (**Fig. 1a**). This was confirmed by the gene expression-based TME deconvolution tool MCPcounter and by immunohistochemical staining with species-specific antibodies (**Fig. 1b-d**, **Extended Data Fig 1a**).

**Figure 1.**
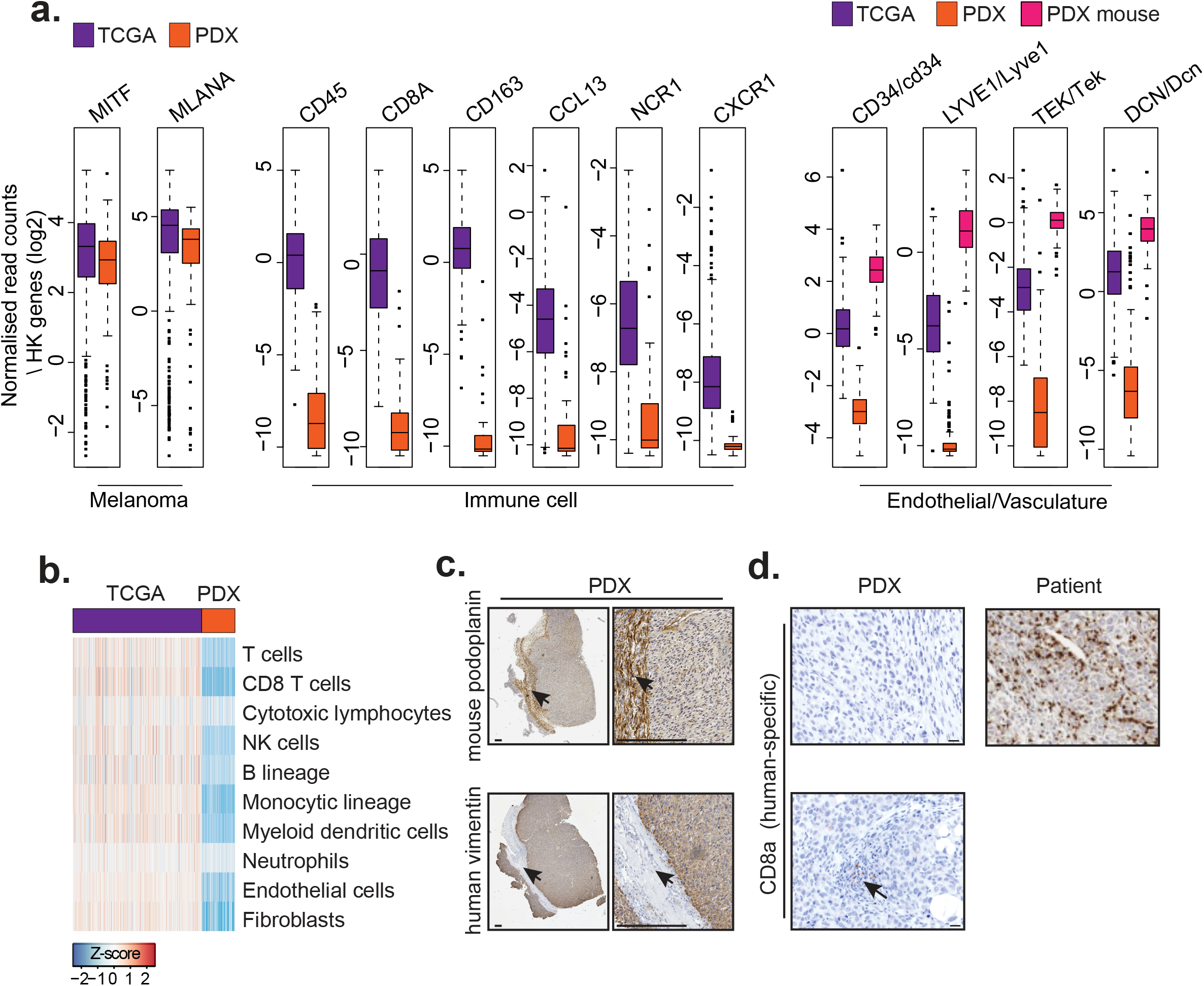
Human TME cells are replaced by mouse cells upon PDX grafting. **a.** Expression of human melanoma genes and TME (immune and vasculature) genes were plotted for patient melanomas from the TCGA database (n=367; purple) and PDX (n=95; orange). For the vasculature genes, also the PDX-carrying mouse gene expression was included (n=95; pink). Gene expression was normalized to the same set of housekeeping genes within each sample. **b.** Loss of human TME gene sets as illustrated by MCPcounter Z-scores, for melanomas in the TCGA (purple) and melanoma PDX (orange). **c.** Samples of PDX M054R.X1 were stained for mouse-specific podoplanin and for human-specific vimentin. Scalebar = 100uM. Arrows indicate the mouse TME cells. **d.** IHC staining of human-specific CD8. Shown are PDX with no cells (top) or sporadic cells positive for CD8 immunostaining (indicated with an arrow), and as comparison a human melanoma.

This observation allowed us to use an algorithm we recently developed (XenofilteR^11^) to distinguish RNA sequence reads of mouse (TME) and human (tumor cell) origin (**Extended Data Table. 2**). In this way, we obtained human-specific tumor cell-intrinsic gene expression values from PDX. These curated signals allowed us to subsequently computationally subtract tumor cell-intrinsic gene expression signals from bulk tumor (that is, tumor cell + TME) gene expression signals from patients in order to establish a TME gene expression signature (**Fig. 2a**). This yielded a set of 767 curated genes expressed in patients’ bulk melanomas, which were (almost) lacking from PDX (**Fig. 2b**, **Extended Data Table 3**). Single cell sequencing (scSeq) data^12^ confirmed that these genes are predominantly expressed by non-tumor cells such as T, B, NK and endothelial cells (**Fig. 2c**, **Extended Data Fig. 1b)**. These data show that by applying the XenofilteR algorithm to PDX data, the RNA signals from tumor cells and TME can be computationally dissected.

**Figure 2.**
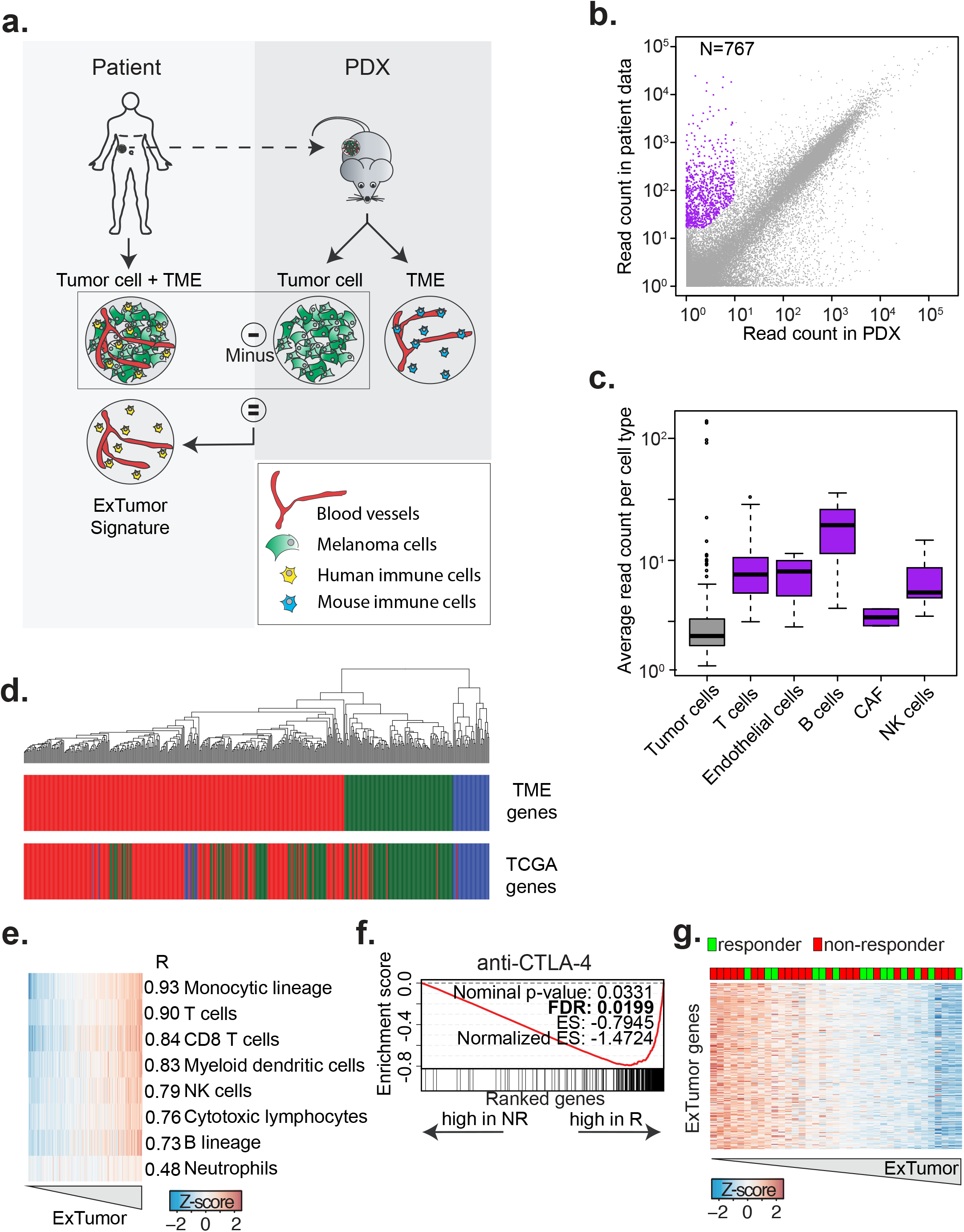
Computational dissection of tumor and TME signals from human melanomas reveals an immune infiltration signature predicting response to anti-CTLA-4 monotherapy. **a.** Graphical representation of the workflow to obtain human tumor cell-intrinsic gene expression values from PDX and the computational subtraction form patient bulk tumor data. Tumor cells are indicated in green, blood vessels in red, human immune cells in yellow and mouse immune cells in blue. **b.** Scatterplot of the average normalized read counts per gene (log10) for human tumor cell-intrinsic gene expression values from PDX (n = 95 samples) and patient samples^9^ (n=133 samples). The identified 767 TME-specific genes are indicated in purple. **c.** Boxplots showing the expression of all 767 TME-specific genes in the tumor cells and 5 TME cell types (in purple) from a single tumor analyzed by single cell sequencing^12^. The median in the boxplots are indicated by the center line, the upper and lower quartiles by the box limits, and the 1.5x interquartile range by the whiskers and outliers are indicated as points. CAF = Cancer Associated Fibroblast. **d.** Cluster analysis of the melanoma TCGA samples (Skin Cutaneous Melanoma, SKCM, N=367) based on the 767 TME-specific genes. Colors indicate the subgroups based on TCGA classification and based on the 767 TME genes. **e.** Pearson correlation of the average expression of the immune-related genes from the TME gene set with specific immune cell types based on melanoma TCGA (SKCM) gene expression data. Expression of the specific cell types was estimated using MCPcounter^26^. **f.** GSEA for the ExTumor signature on patient tumors^16^ before start of treatment with anti-CTLA-4. Genes were sorted according to the expression differences between responders and non-responders to anti-CTLA-4 therapy. Genes on the left side of the plot have a low expression in responders. The genes on the right side have a high expression in responders. **g.** Heatmap with ExTumor signature genes of samples sorted according to response anti-CTLA-4. Patients that responded to anti-CTLA-4 (CR, PR and SD) are indicated in green, non-responders (PD) in red.

## TME signals recapitulate TCGA melanoma classification

In the TCGA, patient melanomas have been classified based on three genetic signatures (“MITF-low”, “Keratin” and “Immune”)^13^. To determine the contribution to this classification of the 767 TME genes that we identified above, we performed a cluster analysis on the TCGA melanoma samples using these TME and immune-related genes only. This revealed a remarkably high overlap with the three melanoma subgroups that had previously been classified using bulk tumor signals (**Fig. 2d**). Cluster analysis with the TME genes in a cohort of 57 samples from stage IV melanoma patients^14^ recapitulated these findings (**Extended Data Fig. 1c**). Thus, by computationally subtracting tumor cell-intrinsic gene expression signals from those in bulk tumor, we uncovered hundreds of TME-specific gene signals, which were sufficient to recapitulate the classification of melanoma subgroups in two independent large tumor datasets. These results illustrate the importance of the signals from the TME, demonstrating that the TCGA classification of melanomas is largely based on gene expression signals that are derived from the tumor signals in the TME, rather than the tumor cell-intrinsic ones.

## Tumor cell-extrinsic signature predicts response to anti-CTLA-4 treatment

The main goal of this study was to generate a predictive classifier for combination ICB treatment. Previously, ICB responses were shown to be affected by both tumor cell-intrinsic and TME factors^6,7^, arguing that a predictive classifier should recapitulate both distinct features. When inspecting the 767 TME genes, we noted that 361 genes showed a high degree of co-regulation (**Extended Data Fig. 2a-b**) and were immune-related (**Ext. Fig. 2c**). The expression of these 361 genes was highly correlated with that of multiple immune cell types, including T-cells (R=0.9) and B-cells (R=0.7) (**Fig. 2e**). With these immune-related genes, which is a measure for the abundance of immune cells in the sample^15^, we built a so-called “ExTumor signature” (**Extended Data Table 4**) and determined whether it is capable of predicting clinical outcome of melanoma patients treated with ICB, whether anti-PD-1 or anti-CTLA-4.

We applied Gene Set Enrichment Analysis (GSEA) on gene expression data from a cohort of patient samples obtained before the start of treatment with anti-PD-1 monotherapy^9^. The ExTumor signature failed to show a correlation with the clinical outcome of this patient cohort (**Extended Data Fig. 2d**). In contrast, when tested on tumors prior to receiving anti-CTLA-4 monotherapy, we observed a significant association between high expression of ExTumor signature genes and clinical response (FDR <0.02, **Fig. 2f-g**) but not with the tumor mutational burden (TMB, **Extended Data Fig. 2e**) This association was validated in an independent cohort of patient samples before start of treatment with anti-CTLA-4^16^ (**Extended Data Fig. 2f**). Again, the ExTumor signature was associated with clinical outcome, showing significantly higher expression in the patients with clinical benefit (p < 0.01, **Extended Data Fig. 2g**) and a better overall survival for those patients with the highest expression (highest quartile) of the ExTumor signature (LogRank p = 0.028, **Extended Data Fig. 2h)**. Thus, the ExTumor signature is strongly correlated with response to anti-CTLA-4, but not anti-PD-1 treatment.

## Tumor cell-intrinsic signature predicts response to anti-PD-1 treatment

Given that the ExTumor signature failed to predict response to anti-PD-1 therapy, we investigated whether, instead, this response could be predicted by tumor-cell intrinsic factors. We again used the curated PDX signals (**Fig. 3a**) and applied a Principal Component Analysis (PCA, which identifies gene sets that explain the largest variance) of the tumor-cell intrinsic RNA expression data (**Extended Data Fig. 3a**). One Principal Component (PC) was associated with anti-PD-1 response (FDR = 0.097, **Fig. 3b** and was used to build a so-called “InTumor” gene expression signature (**Extended Data Fig. 3b and Extended Data Table 4**). We confirmed that the InTumor signature was specific for tumor cell (rather than TME) genes, as shown by *(i)* its high concordance between PDX and matched PDX cell lines (**Extended Data Fig. 4a-b**; *(ii)* IHC with species-specific antibodies (**Extended Data Fig. 3b-g**); and *(iii)* lack of a significant correlation between the InTumor signature expression and immune cells (**Extended Data Fig. 3h**) when using the TCGA database.

**Figure 3.**
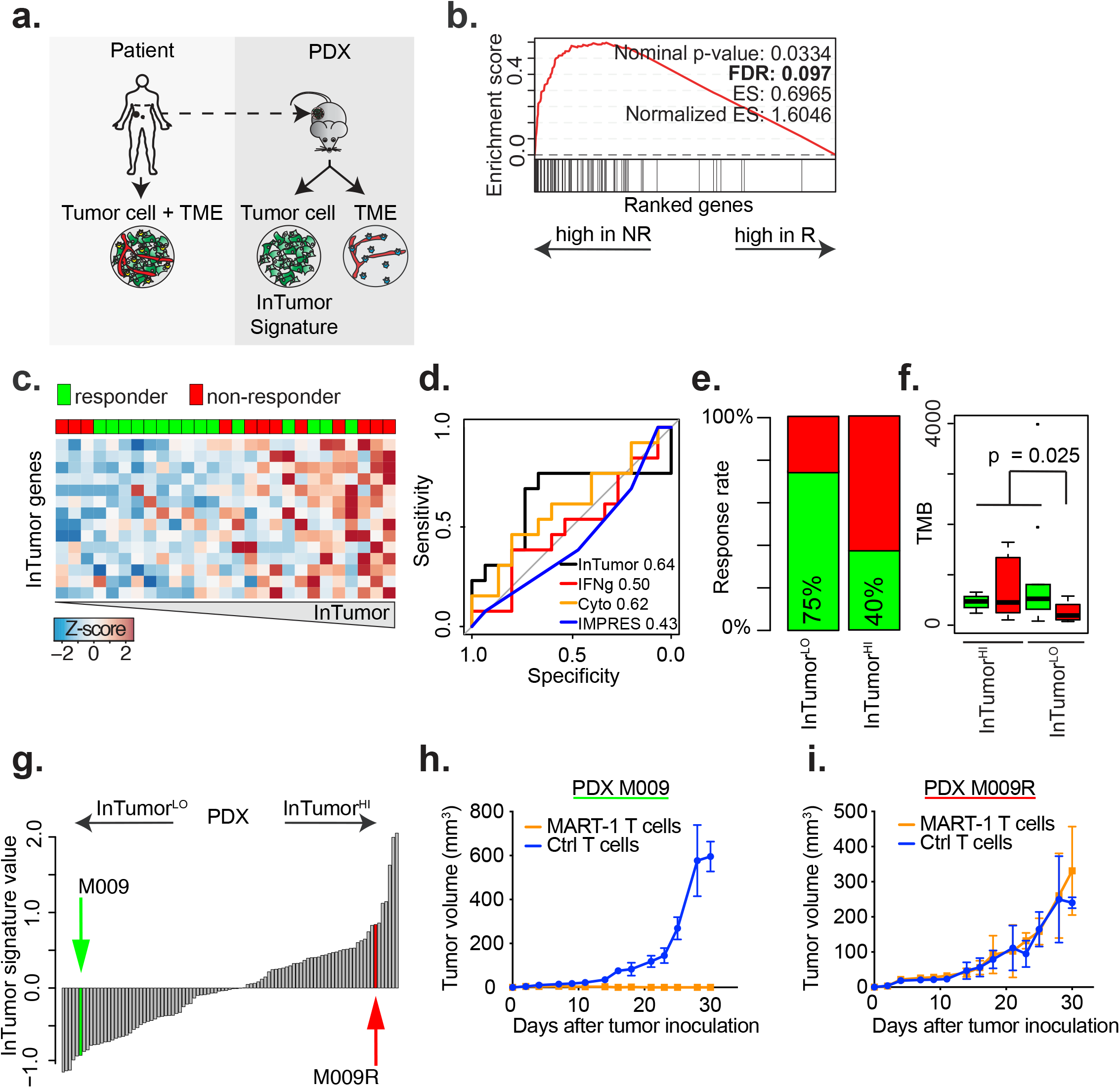
A tumor cell-intrinsic gene expression signature predicts *in vivo* response to T cell killing and to anti-PD-1. **a.** Graphical representation of the PDX platform (n=95 samples) and the acquisition of the curated tumor cell-intrinsic gene expression signals. **b.** GSEA for the InTumor signature on patient tumors^9^ before start of treatment with anti-PD-1.R = Responder (CR, PR and SD), NR = NonResponder (PD) **c.** Expression levels of the InTumor genes in patients who responded to anti-PD-1 treatment and those who did not respond. Samples are ordered according to the average expression of the InTumor signature. **d.** ROC curves for the InTumor, INFγ-signature, Cytolytic score and the IMPRES signature in the patient cohort. The Area Under the Curve (AUC) is indicated after the signature name. **e.** Patients’ response rates to anti-PD-1 separated into InTumor^HI^ and InTumor^LO^ groups. Classification into the two groups (responders in green, non-responders in red) is based on the median InTumor signature value. **f.** Tumor mutational burden (TMB) for responders (green) and non-responders (red) split into the InTumor^HI^ and InTumor^LO^ groups. Pearson’s Chi-squared test with simulated p-value (based on 1×10^5^ replicates). **g.** PDX samples (n=95) ranked according to the average expression of the InTumor signature. Samples from a single patient, but two different time points, are indicated in green (M009) and red (M009R) and were used for in-vivo testing. **h.** Growth curve for M009 PDX sample (passage 6). Mice were either treated with MART-1-specific T-cells (day 1) recognizing tumor (orange) or with Ctr T-cells (blue). **i.** Growth curve for M009R PDX samples (passage 4). Mice were either treated with MART-1-specific T-cells (day 1) recognizing tumor (orange) or with Ctr T-cells (blue).

We next determined whether the InTumor signature predicts response to anti-PD-1. To preclude any effect from non-tumor cells (which are abundantly present in bulk patient tumor sequence data), we first reduced the number of genes of this signature to the 14 showing the highest ratio in expression between tumor and TME (**Extended Data Fig. 5a).** We then performed GSEA on gene expression data from a cohort of patient samples obtained prior to anti-PD-1 monotherapy^9^. This analysis revealed a significantly larger number of non-responding patients in the InTumor^HI^ group (Chi-square p < 0.03; **Fig. 3c**). Furthermore, the InTumor signature outperformed three previous signatures (IMPRES^8^, IFNg^17^ and cytolytic score^18^, **Fig. 3d**). Splitting the samples into two groups based on the median expression resulted in a response rate of 75% in the InTumor^LO^ patients and 40% in the InTumor^HI^ patients (**Fig. 3e**). The predictions were significant, yet imperfect. This could be partly explained by the notion that misclassified predicted responders showed a significantly lower TMB (**Fig. 3f**), in keeping with the association between TMB and ICB response^19^. These results were reproduced in a second cohort^20^ of patients treated with anti-PD-1 mono-therapy (**Extended Data Fig. 5c-f**), demonstrating the predictive power of the InTumor signature. No differences in Tumor Mutational Burden (TMB) were observed between the InTumor^HI^ and InTumor^LO^ groups for either dataset (**Extended Data Fig. 5g**). No link was observed between the InTumor signature and response to anti-CTLA-4 (**Extended Data Fig. 5b**). Thus, the InTumor signature is shows a strong inverse correlation with response to anti-PD-1, but not anti-CTLA-4.

To biologically validate the inverse correlation between the InTumor signature and immune sensitivity, we determined whether this signature predicts T cell killing in a well-defined *in vivo* setting that we developed recently^21^. We selected two melanoma PDX derived from biopsies from the same patient taken before and after progression on vemurafenib (M009 and M009R^10^), which showed a low and high InTumor signature, respectively (**Fig. 3g**). These PDX were transplanted into mice and subsequently challenged with Adoptive T-Cell Transfer (ACT) of TCR-matched cytotoxic T cells^21^. This setup allowed us to challenge these InTumor^LO^ and InTumor^HI^ tumors under identical TME and immune infiltrate conditions. Complete T cell-mediated tumor control was observed for the InTumor^LO^ PDX (M009; **Fig. 3h**). In contrast, the InTumor^HI^ PDX (M009R) expanded exponentially and indistinguishably from the control ACT (**Fig. 3h**, **Extended Data Fig. 6a**). These results were confirmed in four melanoma cell lines with different InTumor scores (**Ext Fig. 6b-c**). No correlation between response and the infiltration of T cells was observed (**Extended Data Fig. 6a, d**). These results confirm that an InTumor^HI^ signature is associated with resistance to T cell killing *in vivo*.

## InTumor and ExTumor signatures jointly predict response to anti-CTLA-4 or anti-PD-1 combination treatment

The establishment of the ExTumor signature predicting anti-CTLA-4 response and the InTumor signature predicting anti-PD-1 response allowed us to ask next whether these signatures, alone, had predictive value for the clinical response to either anti-CTLA-4 or anti-PD-1 in the combination treatment setting. This was determined for stage IV melanoma patients (CA209-038 trial, NCT0162149023^20,22^). ROC analysis revealed that both individual signatures had predictive value, but underestimated the number of patients responding to combination ICB (**Extended Data Fig. 7a-d**).

Since the ExTumor and InTumor signatures independently underestimated the number of responders to combination ICB, we next determined whether they, when used together, have better predictive power for combination ICB. Indeed, the signatures when used in conjunction correctly predicted outcome for combined ICB for 5/6 patients who had relapsed and 14/15 patients who remained relapse-free (chi-square p=0.003, AUC = 0.92, **Fig. 4a**). The sensitivity and specificity of this ICB response quadrant model outperformed previous signatures (IMPRES^8^, IFNg^17^ and cytolytic score^18^, **Extended Data Fig. 7e)**. Furthermore, the combined signature prediction showed a significant difference in progression-free survival (PFS) between predicted combination ICB responders and non-responders (logrank, p=0.0001, **Fig. 4b**). Importantly, we confirmed these results in the independent OpACIN trial^23^ (NCT02437279) in which stage III melanoma patients were treated also with anti-PD-1 + anti-CTLA-4. Again, the ExTumor and InTumor signatures used in conjunction correctly predicted outcome for combined ICB for 4/6 non-responders and 11/12 responders (chi-square p=0.04, AUC = 0.792, **Extended Data Fig. 7f-g**) which was independent of tumor mutational burden (**Extended Data Fig. 7h**). Furthermore, the combined signature prediction showed a significant difference in progression-free survival (PFS; logrank, p=0.0007) between predicted responders and non-responders (**Extended Data Fig. 7i**).

**Figure 4.**
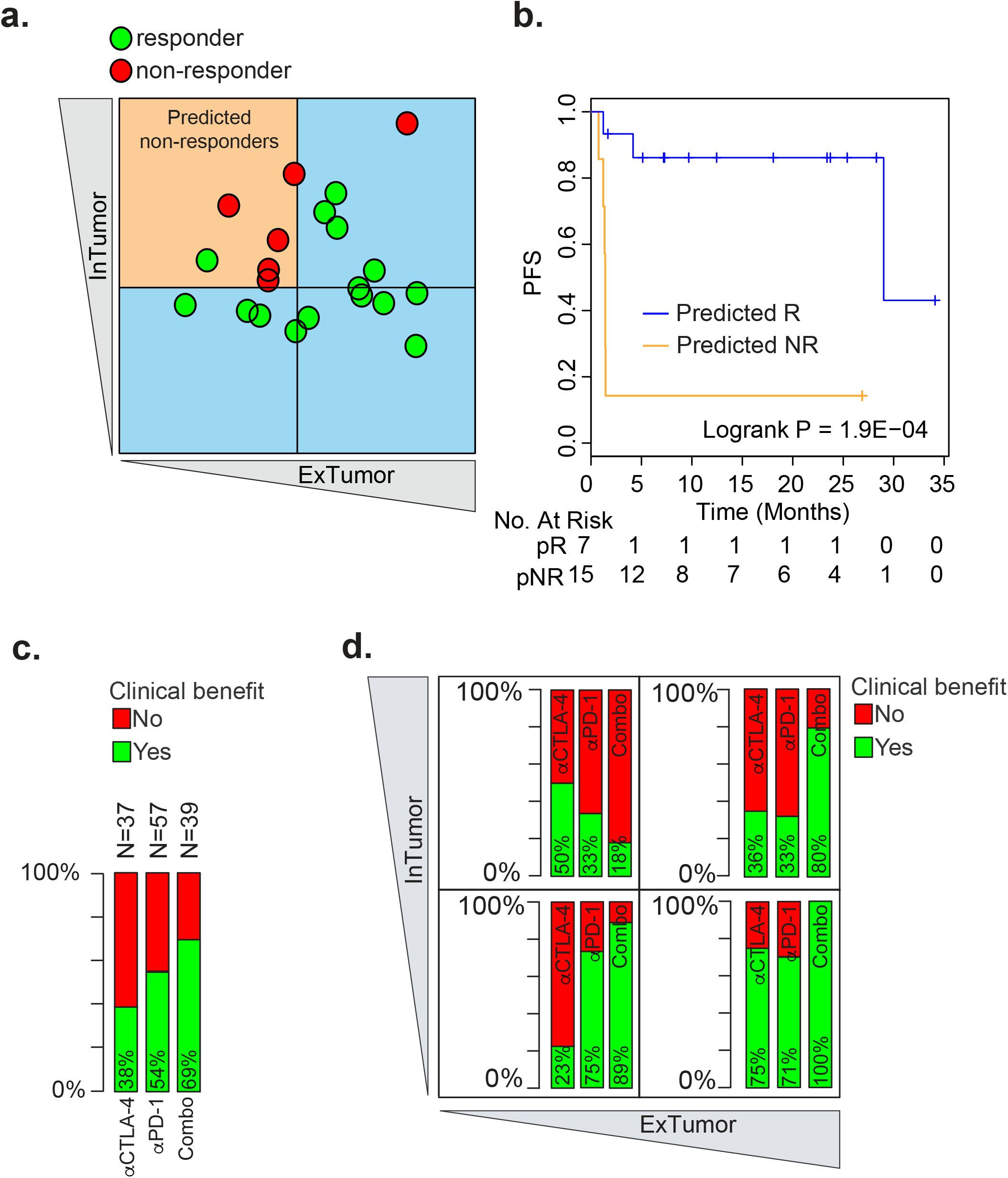
Joint InTumor and ExTumor signatures predict response to ICB combination therapy. **a.** The InTumor and ExTumor signatures used in conjunction on the samples from the CA209-038 clinical trial. Green circles indicate patients who responded to combination ICB therapy, the red circles indicate patients who did not respond to combination ICB. Response is assessed based on the best overall response (RECIST1.1). The three blue quadrants indicate InTumor and ExTumor signatures value for samples that are predicted as responders to either anti-CTLA-4, anti-PD-1, or both. The orange quadrant indicates the InTumor and ExTumor signature values for samples that are predicted as non-responders to anti-CTLA-4 as well as anti-PD-1. **b.** Kaplan-Meier plot comparing recurrence-free survival (PFS) for patients predicted as responders (pR; blue) to patients predicted as non-responders (pNR; orange) for the CA209-038 trial. **c.** Bar graph showing the percentage of patients with clinical benefit to anti-CTLA-4 (left bar, van Allen et al. data^16^, n=37), anti-PD-1 (middle bar, Hugo et al.^9^ + Riaz et al.^20^ data, n=57) and anti-CTLA-4 + anti-PD-1 combination therapy (right bar, Blank et al.^23^ + CA209-038 data, n=39). **d.** Quadrant plot based on the InTumor and ExTumor signatures illustrating the clinical benefit to ICB in the four groups based on the expression of the InTumor and ExTumor signatures. Bar graph shows the percentage of patients with a clinical benefit to anti-CTLA-4 (left bar, van Allen et al. data, n=37), anti-PD-1 (middle bar, Hugo et al. + Riaz et al. data, n=57) and anti-CTLA-4 + anti-PD-1 combination therapy (right bar, Blank et al. + −038 data, n=39).

Lastly, we combined the InTumor with the ExTumor signatures to determine the percentage of responders to monotherapy (either anti-PD-1^9,20^ or anti-CTLA-4^16^) and combination ICB therapy (anti-PD-1 + anti-CTLA-4^22,23^) by classifying them into so-called response quadrants (5 cohorts, n=131 patients, **Fig. 4C**). Both for anti-PD-1 mono-therapy and the combination therapy, the classification of patients in these cohorts into the quadrants was significantly associated with clinical benefit (Chi-square, p < 0.001 and p < 0.05, resp. **Fig. 4c**). More importantly, this analysis revealed that patients with an InTumor^HI^/ExTumor^HI^ classification respond significantly better to combination (anti-CTLA-4 + anti-PD1) treatment (80% of patients had clinical benefit) than to either anti-CTLA-4 or anti-PD-1 alone (36% and 33% resp., Chi-square p < 0.05, **Fig. 4d**, **Extended Data Fig. 8a).** Furthermore, the patients classified as InTumor^LO^/ExTumor^LO^ did not significantly benefit from anti-CTLA-4 in addition to anti-PD-1 (Chi-square p < 0.05, **Fig. 4d**, **Extended Data Fig. 8b**), demonstrating that this ICB response quadrant model has the potential to perform personalized response prediction and thereby guide treatment choice.

CTLA-4 and PD-1 blockade have become standard checkpoint inhibitor therapies in various malignancies including melanoma, bladder cancer and renal cancer. These antibodies are administered either as mono- or combination therapy. However, because the toxicity profile of the combination is significantly worse, clinicians need tools to decide whether PD-1 monotherapy is likely to give sufficient clinical benefit, or whether combination with CTLA-4 blockade is required for improved clinical outcome. By identifying tumor cell-derived “InTumor” and tumor cell-extrinsic “ExTumor” gene expression profiles, we were able to develop a classification for individual patients into four treatment quadrants, each covering a distinct response to both ICB mono-therapies and the ICB combination therapy. Biologically, we demonstrate that melanomas classified with a high InTumor signature score are refractory to T cell elimination in mice. However, similar to several other types of gene expression signatures^8,9,24^, whether the classifiers contain one or more genes that mechanistically contribute to these biological and immunological traits will need further study. A possible candidate might be NGFR, one of the 68 genes in the InTumor signature. We recently showed that it contributes to resistance to multiple therapies including ICB^25^. With the validation of these classifiers both in animal models and in several independent patient cohorts, we demonstrate their clinical relevance as critical biomarkers for single and combination ICB treatment response. Our results will need to be validated in additional larger patient cohorts before they may become a decision tool in the daily clinical practice. These signatures may enable personalized treatment advice based on the predicted clinical benefit from either anti-CTLA-4 or anti-PD-1 monotherapy or from their combination, or point to the need for alternative checkpoint inhibitor combination therapy.

## Acknowledgements

We would like to express our gratitude to the patients and their relatives for their cooperation in this study. We acknowledge NKI-AVL Animal Pathology facility for performing the immunohistochemistry stainings and the NKI Research-HPC facility for computational resources. We would like to thank the Peeper and Blank research groups, Lodewyk Wessels and Karin de Visser for valuable input, Rachid Manumur for bioinformatical analysis and Steven de Jong for the vimentin IHC protocol. The research leading to these results has received funding from the European Research Council under the European Union’s Seventh Framework Programme (FP7/2007-2013) / ERC synergy grant agreement n° 319661 COMBATCANCER. This work was financially supported by a grant from the Dutch Cancer Society (NKI-2013-5799; K.K., P.C.-S. and D.S.P.), Cancer Research UK and The Wellcome Trust (WT098051, D.J.A.), and a Queen Wilhelmina Award by the Dutch Cancer Society (D.S.P.).

