## Supplementary methods and figures for "PREDICTIVE IMMUNE CHECKPOINT BLOCKADE CLASSIFIERS DISTINGUISHING MONO- VERSUS COMBINATION THERAPY REQUIREMENT"

#### PDX and PDX-derived cell line generation

The collection and use of human tissue were approved by the Medical Ethical Review Board of the Antoni van Leeuwenhoek. Animal experiments were approved by the animal experimental committee of the institute and performed according to Dutch law. PDX were generated as described in Kemper et al.[1]. PDX-derived cell lines were generated after enzymatic dissociation (described in1) by growing the cells in 2D in 10% fetal bovine serum (FCS, Sigma), 2 mM glutamine, 100 U/ml penicillin, and 0.1 mg/ml streptomycin (all Gibco) under standard conditions. A full report of PDX relevant information following the guidelines described in Minimal Information for Patient-Derived Tumor Xenograft Models (PDX-MI) is available (**Extended Data Table 1**)[2].

#### Verification of origin of PDX and PDX-derived cell lines

STR profiling was performed on gDNA from all PDX and PDX-derived cell lines (**Extended Data Table 1**). gDNA was isolated by using DNA Easy Blood & Tissue Kit (Qiagen) according to manufacturer's protocol. STR profiling was performed using the Powerplex 16 HS kit (DC2101, Promega) according to manufacturer's protocol. The samples were run on an ABI3700 and analyzed using GeneMapper. In addition to the STR profiling we performed a SNP analysis on both the RNA sequence data from the PDX cell lines as well as on RNA sequence data of the first passage PDX. The percentage of overlap in SNPs was tested between all cell lines, between all first passage PDX samples and between PDX and PDX cell lines which confirmed PDX cell lines origin (**Extended Data Fig 9**).

**Extended Data Table 2: STR profiling of PDX-derived cell lines.**

| SampleID | AMEL | CSF1PO | D13S317 | D16S539 | D5S818 | D7S820 | TH01 | TPOX | vWA |
|---|---|---|---|---|---|---|---|---|---|
| M002.X1.CL | X | 10,12 | 12,14 | 12 | 12 | 9,10 | 6 | 8,9 | 15,18 |
| M016.X1.CL | X,Y | 12 | 11,13 | 11,12 | 12 | 8,9 | 7,8 | 8 | 14,18 |
| M019R.X1.CL | 11,12 | 11 | 11,15 | 12 | 9,11 | 6 | 11 | 16 | |
| M026.X1.CL | X,Y | 9,11 | 10,13 | 13 | 11,12,15 | 10,11 | 7,9 | 8,9 | 16,17 |
| M026R.X1.CL | X,Y | 9,11 | 10,13 | 13 | 11,12,15 | 10,11 | 7,9 | 8,9 | 16,17 |
| M027.X1.CL | X | 11,12 | 11,12 | 11,12 | 11,12 | 11 | 6,9 | 8,11 | 17,19 |
| M029.X1.CL | X | 14 | 11,12 | 9,13 | 10 | 11 | 9.3 | 8,11 | 16 |
| M029R.X1.CL | X | 14 | 11,12 | 9,13 | 10 | 11 | 9.3 | 8,11 | 16 |
| M032.X2.CL | X,Y | 10 | 11,12 | 9,12 | 12,16 | 12,13 | 7 | 8,11 | 17,18 |
| M032R1.X1.CL | X,Y | 10 | 11,12 | 9,12 | 12,16 | 12,13 | 7 | 8,11 | 17,18 |
| M032R5.X1.CL | X,Y | 10 | 11,12 | 9,12 | 12,16 | 12,13 | 7 | 8,11 | 17,18 |
| M032R6.X1.CL | X,Y | 10 | 11,12 | 9,12 | 12,16 | 12,13 | 7 | 8,11 | 17,18 |
| M036-2.X1.CL | X,Y | 8 | 9,12 | 8 | 12 | 12 | 9,9.3 | 8,11 | 16,18 |
| M044R.X1.CL | X | | 11 | 8,13 | 11 | 8 | 6,9.3 | 8 | 19 |
| M048R.X1.CL | X | 11 | 11,14 | 10,11 | 12 | 12 | 6 | 8 | 16,19 |
| M050.X2.CL | X | NA | 12,13 | 11 | 13 | 8,11 | 7,11 | 8,11 | 16 |
| M061R.X1.CL | X,Y | 10 | 12,14 | 9 | 10 | 9,10 | 6 | 11 | 17 |
| M063R.X1.CL | X | 10,11 | 9,14 | 9,12 | 13 | 8,11 | 8,9 | 8,11 | 17,19 |
| M074R.X1.CL | X | 8 | 11,14 | 8,9 | 12,13 | 11 | 6,9.3 | 8,11 | 15,18 |
| M080.X1.CL | X | 7,9 | 8,11 | 12 | 11 | 10,11 | 9.3 | 8 | 18,19 |
| M082.X1.CL | X,Y | 8,10 | 11 | 8,9,11 | 12 | 10,12 | 7 | 8,9 | 17 |
| M091.X1.CL | X,Y | 10 | 12 | 10 | 11,13 | 9,10 | 7,9 | 8 | 16,17 |

**RNA isolation and sequencing and mapping to mouse and human genomes**
RNA was isolated from PDX and PDX-derived cell lines using Trizol according to manufacturer's protocol. Next, samples were cleaned twice by using the RNeasy MinElute Cleanup Kit (74204, Qiagen) according to manufacturer's protocol, before they were submitted for quality control by using the Agilent 2100 Bioanalyzer system according to manufacturer's protocol. A total of 95 (PDX) and 22 (PDX-derived cell lines) samples were sequenced in 2 batches. PDX samples were uniquely barcoded and pooled into a single stranded library and send for sequencing. To minimize the possible lane effect, we adopted the strategy of pooling all uniquely labelled PDX samples and running the pool on multiple lanes on Illumina HiSeq 2000 sequencers. An average 56 Million unique read pairs were sequenced per sample ranging from 33M to 78M. The same pooled strategy was used for the 22 PDX cell lines; only these samples were sequenced to an average of 39 Million unique reads pairs, ranging from 27M to 54M. All samples were mapped with Tophat2 v2.1.0[3] with the following parameters: --library-type fr-firststrand -g 1 -p 8 -G ENSEMBL_Annotation_v82.gtf. All samples were mapped to human reference, GRCh38 (ENSEMBL v82) and mouse reference genome GRCm38 (ENSEMBL v82).

**Computational dissection of sequence reads from human and mouse origin**
Sequence data from PDX samples are contaminated with sequence reads of mouse origin[4]. Therefore, computational removal of these sequence reads is pivotal. Previously we developed XenofilteR which showed accurate filtering, both for RNAseq and DNAseq data. To further test the performance of XenofilteR in our RNAseq samples we mixed the sequence reads from a mouse cell line (~25%) with the sequence reads from a human cell line (~75%). Both the original mouse, human and mixed sequence reads were mapped to the human and mouse reference genome. The resulting bam file from the mixed sample was subsequently filtered using XenofilteR (version 1.4, github.com/PeeperLab/XenofilteR) with default settings. Furthermore, the RNA isolation protocols and sequencing for these 2 cell lines were identical to that of the PDX RNA sequencing described here. Filtering with XenofilteR retained 99.14% of human sequence reads while removing 99.7% of mouse reads (**Extended Data Table 2**). The removal of mouse sequence reads in the mixed sample resulted in read count data similar to the read counts when only the human sequence reads were mapped. Thus, XenofilteR shows accurate filtering of mouse sequence reads while retaining sequence reads of human origin.

**Extended Data Table 2: Number of sequence reads for the human cell line, the mouse cell lines and the mixed sample before and after filtering with XenofilteR.**

| Sample | Human reads | Mouse reads | Total reads | % Mouse reads |
|---|---|---|---|---|
| **Human** | 46172045 | 0 | 46172045 | 0.0% |
| **Mouse** | 0 | 14082189 | 14082189 | 100.0% |
| **Mixed_unfiltered** | 46170355 | 14081047 | 60251402 | 23.4% |
| **Mixed_filtered** | 45814444 | 121160 | 45935604 | 0.3% |

Next, we applied XenofilteR[4] to the 95 PDX samples. The percentage of reads from mouse origin in each PDX RNA sequencing sample was measured by counting the unique read pairs in the bam files with mouse specific reads as generated with XenofilteR divided by the total number of unique read pairs (sum of unique read pairs mapped to either mouse or human after filtering with XenofilteR).

**Gene expression analysis**

Bam files filtered from sequence reads of mouse origin, and thus retaining only read of human origin. To generate the curated read count data bam files were name sorted with picard followed by counting reads with HTseq-count (HTSeq-0.6.1p1)[5] with settings: -m intersection-nonempty -a 10 -i gene_id -s reverse -f bam. Count data generated with HTseq-count was analyzed with DESeq2[6]. Centering of the normalised gene expression data is performed by subtracting the row means and scaling by dividing the columns by the standard deviation (SD) to generate a z-score.

**ExTumor signature**

The curated tumor cell-intrinsic read count data of 95 PDX samples were used to identify the gene expression signals from the human tumor microenvironment (TME) in total patient tumors (tumor + TME). For this we used a dataset of 133 patient melanoma samples[7] for which pre-processing was performed identical as for the PDX samples. To identify the human TME specific signals from the patient samples we selected the genes with a minimal read count of 20 and with at least a log2-fold difference of 8 in expression between curated tumor cell-intrinsic signals (tumor) from PDX and total patient tumors (tumor + TME). This yielded 787 TME specific genes. To verify these TME specific signals we tested the expression in single cell RNA sequence data (scSeq) downloaded from the NCBI GEO database (GSE72056)[8]. Count data from each cell was normalized to 1 million reads per cell. Classification into somatic and germ-line and specific cell types were taken from the original publication6. The expression of the 787 TME specific genes was tested in the scSeq data from the three patients (sample ID 79, 80 and 88, **Fig. 2c & Extended Data Fig. 1b**) with the highest number of cells sequenced for tumor cells as well as TME cells (T-cells, Endothelial cells, B-cells, CAFs and NK-cells).

To further specify the 787 TME specific signals we performed cluster analysis using the TCGA SKCM dataset (**Extended Data Fig. 2a**). For this, raw read count data was downloaded using “TCGAbiolinks” package[9] and pre-processed and normalized similar to the PDX dataset. Cluster analysis was performed using the metastatic melanoma samples from the TCGA database[10] and the 767 genes (**Extended Data Fig. 2b**). This analysis revealed 2 clusters with highly co-regulated genes. The first cluster was dominated by keratin genes and signals that origin from keratinocytes, the second cluster (361 genes) was highly enriched for immune cell related signaling. The latter was confirmed by Gene Ontology analysis (**Extended Data Fig. 2c**) showing the GO-pathways ‘inflammatory response’ and ‘Immune response’ as top hits. These 361 genes comprise the ExTumor signature (**Extended Table. 2**).

**InTumor signature**

The curated tumor cell-intrinsic read count data of 95 PDX samples[1] were used to perform a principle component analysis (PCA) to identify the signals that explain the largest variation in the dataset (**Extended Data Fig. 3a**). The first three principle components (PC) together explained 22.5% of the variance in the dataset (11.3%, 6.9% and 4.4%, respectively). These three PC’s were used to generate three gene expression signatures by selecting the genes for each of the three principle components with the highest rotation value and comprise 39, 89 and 76 genes respectively (**Extended Data Table. 5, Extended Data Fig. 3b**). Next, we performed gene set enrichment analysis (GSEA) to correlate the expression of the three gene sets with response to immune checkpoint blockade in two datasets (mono-therapy anti-PD-1[11] and mono-therapy anti-CTLA-4[12]). Only significantly association between the PCs and response to anti-PD-1 was the second PC with response to anti-PD-1. The geneset generated using PC2 comprises 89 genes and is named the InTumor signature.

The InTumor signature was developed on PDX samples without signals from the TME. However, this does not preclude the possibility that these genes are expressed also in stromal cells. To rule out any possible influence from signals of TME origin in patient samples we checked the average expression of the signature genes in both tumor and stromal cells using scSEQ data. For each gene in the InTumor signature we calculated the average expression in tumor cells and the stromal cells using scSEQ data[8]. Samples were normalised to 1M reads per samples and the three samples with high number of tumor as well as stromal cells were used for the analysis (Samples 79, 80 and 88). Only genes with an arbitrary 10-fold higher expression in tumor cells compared to stromal cells comprise the InTumor exclusive signature (**Extended Data Fig. 5a**). Out of the 86 genes we identified 14 genes with a 10-fold higher expression in tumor compared to stromal cells (**Extended Data Table. 6**). These 14 genes were used to measure the InTumor signature in patient samples.

**ExTumor and InTumor signature values**

Signature values (ExTumor and InTumor) were generated by calculating the average gene expression of the signature genes per sample. This was performed on the z-scores as calculated per dataset. Because the z-score is a median centered expression value over the dataset this will result in an equal distribution of samples in a dataset that are divided into signature$^{HI}$ and signature$^{LO}$ groups. These cut-offs are not trained for and are used identical in all datasets including for the datasets with patients treated with the combination of anti-CTLA-4 + anti-PD-1 and the separation of patients in the ICB response quadrant.

**Validation dataset Nanostring**

Gene expression of 51 samples was assessed using the nCountera PanCancer Immune Profiling assay by NanoString Technologies (Seattle, WA, USA). RNA was isolated from formalin fixed, paraffin embedded tumor samples using the QIAgen AllPrep DNA/RNA kit on the QIAcube according using standard manufacturer's protocol. Pre-processing and data normalisation were performed using the nSolver (Nanostring) software using default settings. The ExTumor signature value was calculated using the 151 genes available on the platform. All samples were taken before start of treatment with anti-CTLA-4. Available clinical variables included survival data for all patients and response to anti-CTLA-4 for 42 out of the 51 patients. Patients with progressive disease (PD) were indicated as non-responders. Patients with stable disease (SD), partial response (PR) and complete response (CR) were indicated as responders.

**Validation datasets RNAseq and Exome sequencing**

For all patient datasets that were used to validate the ExTumor and InTumor signatures raw sequence data (fastq files) was downloaded. All samples were mapped to the human reference genome (Homo.sapiens.GRCh38.v82) using STAR(2.6.0c)[13]

with default settings. Count data generated with HTseq-count was analyzed with DESeq2[6]. Centering of the normalised gene expression data per dataset was performed by subtracting the row means and scaling by dividing the columns by the standard deviation (SD). The generated z-score used to calculate the signature values. For all datasets the identical settings were used except for the OpACIN sequence data as these were sequenced single-end whereas all other datasets were sequence paired-end. For the validation analyses the cutaneous melanomas were used. These classifications were based on the clinical data as reported by each publication.
To assess tumor mutation burden (TMB) for the melanoma patient datasets (van Allen[12], Riaz[14] and OpACIN[15]) raw sequence reads were downloaded (fastq) and mapped to the human reference genome using BWA[16]. Subsequently, the GATK4 pipeline was used to mark duplicate sequence reads and recalibrate base quality scores followed by mutation calling using Mutect2. TMB was calculated by summarizing the number of non-synonymous mutations with a minimum of 15 sequence reads and a variant allele frequency of 5%. For the Hugo datasets raw sequence reads were not available, therefore the TMB was taken as reported in the supplementary data of the Hugo et al. paper[11].

**Comparison of InTumor with published gene expression signatures**

Three previously published gene expression signature that were designed to predict response to ICB were tested and its predictive power compared to the predictive power of the InTumor signature. The cytolytic score[17] was defined as the average expression (Z-Score) of the genes PRF1 and GZMA for each sample. The IFNg[18] score was defined as the average expression (based on the Z-Score) of 10 genes (CXCL11, IDO1, CXCL9, IFNG, PRF1, STAT1, HLA-DRA, CCR5, GZMA and CXCL10). To calculate the IMPRES[19] score we generated FPKM values based on the read count data for each dataset. For each sample the ratio between 15 gene pairs was assessed (“CD274/VSIR", "CD28/CD276", "CD86/TNFRSF4", "CD86/CD200", "CTLA4/TNFRSF4", "PDCD1/TNFRSF4", "CD80/TNFSF9", "CD86/HAVCR2", "CD28/CD86", "CD27/PDCD1", "CD40/CD274", "CD40/CD80", "CD40/CD28", "CD40/CD274", "TNFRSF14/CD86"). The IMPRES score is the sum of the number of pairs where the expression of the first gene is higher than the second. This value will be between 0 and 15.

**Gene Set Enrichment Analysis**

GSEA based on the principle components was performed using the BROAD javaGSEA standalone version (http://www.broadinstitute.org/gsea/downloads.jsp) and the curated ‘hallmark genesets’ (http://software.broadinstitute.org/gsea/msigdb/collections.jsp). Analysis were run using 10,000 permutations. Genes were ranked based on the Signal2Noise metric. Genesets with an FDR <0.1 were considered significant.

**qPCR**

cDNA was generated using either Superscript III Reverse Transcriptase or Maxima First Strand cDNA synthesis kit for RT-PCR according to manufacturer’s protocol. qPCR was performed as described previously[20].

**Extended Data Table 8: Used primers and sequence**

| Gene name | Huma n or mouse | Forward | Reverse |
|---|---|---|---|
| *gapdh* | Mouse + human | GCCAAGGTCATCCATGACAACT | GAGGGGCCATCCACAGTCTT |
| *hprt* | Mouse | CTGGTGAAAAGGACCTCTCG | TGAAGTACTCATTATAGTCAAGGGC A |
| *HPRT* | Human | CGGCTCCGTTATGGCG | GGTCATAACCTGGTTCATCATCAC |
| *ABCB5* | Human | ATTGGAGTGGTTAGTCAAGAGC C | AGTCACATCATCTCGTCCATACT |
| *NGFR* | Human | CCGTTGGATTACACGGTCCAC | TGAAGGCTATGTAGGCCACAA |
| *SPOCK 1* | Human | ACCCCTGCCTGAAGGTAAAAT | GGCTTGCACTTGACCAAATTC |
| *NATL8* | Human | CTACAGCCGCAAGGTGATCC | GAGTCCACAGACATCCGCA |

#### Immunohistochemistry

CD8, PD-L1 and vimentin stainings were performed manually. FFPE slides were deparaffinized and subjected to antigen retrieval using TRIS/EDTA buffer. Slides were blocked in 4% BSA and 4% Normal Goat serum (NGS) in PBS for 30 min, and subsequently incubated overnight at $4^0$C with CD8 (M7103, DakoCytomation), PD-L1 (1795-1-AP, Proteintech) or vimentin antibody (M0725, DAKO) at a dilution of 1:2000 in 1% BSA and 1.25% NGS in PBS. After rinsing with 0.05% Tween-20 in PBS, the slides were incubated with secondary antibody (Goat, Southern Biotech) for 30 min at a dilution of 1:100, and then incubated with streptavidin/HRP (DakoCytomation; P0397) for 30 min at a dilution of 1:200, both in 1% BSA and 1.25% NGS in PBS. Stainings were visualized using 3,3 -diaminobenzidine (DAB) chromophore (Sigma; D-5905) and counterstained with hematoxylin. Sections were reviewed with a Zeiss Axioskop2 Plus microscope (Carl Zeiss Microscopy, Jena, Germany) and images were captured with a Zeiss AxioCam HRc digital camera and processed with AxioVision 4 software (both from Carl Zeiss Vision, München, Germany).

#### Cell lines used for in vivo experiments

All melanoma cell lines used for in vivo ACT models were obtained from the Peeper laboratory cell line stock. PDX were generated as described previously[1]. Cell lines were regularly confirmed to be mycoplasma-free by PCR. Melanoma cell lines were cultured in DMEM (Gibco), with fetal bovine serum (Sigma), 100 U/ml penicillin and 0.1 mg/ml streptomycin (both Gibco) under standard conditions. MART-126-35 and HLA-A2 were introduced in the cell lines (D10, A375, BLM and SkMel-147) using lentiviral constructs. Constructs for lentivirus were packaged using two helper plasmids (psPax and MS2G, Addgene) in HEK293T cells. MART-126-35 -Katushka positive cells were sorted by flow cytometry.

#### Isolation and generation of TCR-specific CD8 T cells

MART-1 (1D3) TCR retrovirus was produced in a packaging cell line as described previously[21]. Peripheral blood mononuclear cells were isolated from healthy donor buffycoats (Sanquin, Amsterdam, the Netherlands) by density gradient centrifugation using Lymphoprep (Stem Cell Technologies). CD8+ T cells were purified using CD8 Dynabeads (Thermo Fisher Scientific), activated for 48 hours on a non-tissue culture treated 24-well plate that was pre-coated overnight with αCD3 and αCD28 antibodies (eBioscience, 16-0037-85 and 16-0289-85) at 2 x 106 per well. Activated CD8 T cells were harvested and mixed with TCR retrovirus and spinfected on a Retronectin coated (Takara, 25µg per well) non-tissue culture treated 24-well plate for 2 hours at 2000g. After 24 hours, T cells were harvested and maintained in RPMI (Gibco) containing 10% human serum (One Lamda), 100 units per ml of penicillin, 100µg per ml of streptomycin, 100 units per ml IL-2 (Proleukin, Novartis), 10ng per ml IL-7 (ImmunoTools) and 10ng per ml IL-15 (ImmunoTools).

**Animal studies**

All animal studies were approved by the animal ethics committee (AEC) of the Netherlands Cancer Institute (NKI) and performed in concordance with ethical and procedural guidelines established by the NKI and Dutch legislation. For xenograft studies, 1 x 106 tumor cells were injected subcutaneously into NSG-□2Mnull mice (Jax). Growth was monitored three times per week with calipers, and tumor size was calculated using the following formula: ½ × length (mm) × width (mm). When tumors reached indicated sizes, mice were randomized over different treatment groups in a blinded fashion and were administered 5 x 106 human CD8 T cells, corrected for transduction efficiency, intravenously in the tail vein. In vivo persistence of T cells was stimulated by administering 100.000 U IL-2 (Proleukin, Novartis) intraperitoneally daily for three consecutive days. All experiments ended for individual mice either when the tumor volume exceeded 1000 mm3, when the tumor showed ulceration, in case of serious clinical illness, when the tumor growth blocked the movement of the mouse, or when tumor growth assessment had been completed.

**Data availability**

RNA sequence data of the 95 PDX samples and 22 PDX derived cell lines have been deposited in the ncbi GEO database under accession number: GSE129127.

**Patient datasets**

TCGA gene expression data was downloaded using the R/Bioconductor package 'TCGAbiolinks'9 as read count values. RNA sequence data from the Hugo et al. papers10,11 were download from NCBI's NCBI's GEO database (GSE78220 & GSE65186). RNA sequence data from the OpACIN trial[15] is available from the EGA database: EGAS00001003099. Read count data from the CA209-038 trial data is available as **Ext. Data Table 7**. Single cell RNA sequence data was downloaded from NCBIs GEO database: GSE72056.

**Code availability**

All pre-processed data and code to recreate the figures from this manuscript will be available through Github (Peeperlab/Krijgsman_etal) upon publication.

#### Extended data legends

**Extended Data Fig. 1. Replacement of human stromal cells by mouse stromal cells**. **a.** IHC staining of PDX sample M005.X1 with mouse podoplanin and human vimentin antibodies, showing replacement of human stromal cells with mouse stromal cells within one passage after grafting in mice. **b**. Expression of the 767 stromal-derived genes using 2 samples from scSeq data (samples 80 and samples 79). **c.** Heatmap and cluster analysis of 57 stage IV melanoma samples based on the 767 stromal-specific genes. The three stromal sample clusters highly overlap with previously identified subgroups which was based on whole tumor signals.

**Extended Data Fig. 2. Development and validation of the ExTumor signature. a.** Flow chart of the development of the ExTumor signature. **b.** Hierarchical clustering the 767 tumor-cell extrinsic genes using the TCGA metastatic melanoma samples. The highly correlated genes in the top right cluster were identified as high in immune-related genes and named the 'ExTumor' signature. **c.** Gene Ontology pathway analysis with the genes identified in a. (ExTumor signature). **d** GSEA for the ExTumor signature on patient tumors before start of treatment with anti-PD-1. Genes were sorted according to the expression differences between responders and non-responders to anti-PD-1 therapy. Genes on the left side of the plot have a low expression in responders. The genes on the right side have a high expression in responders. **e.** Tumor mutational burden (TMB) in $ExTumor^{HI}$ and $ExTumor^{LO}$ samples. **f.** Validation cohort with samples before start of treatment with anti-CTLA-4. Heatmap shows the expression of the genes measured by Nanostring nCounter. The samples ordered according to the ExTumor signature expression. Responders and non-responders to therapy are indicated in green and red, respectively. **g.** Difference in expression of the ExTumor signature between responders and non-responders to anti-CTLA-4 therapy (Pvalue < 0.01, 2-sided Ttest). **h.** MK-plot showing the difference in survival between patients with an $ExTumor^{HI}$ (top 25% of patient samples with highest expression) and an $InTumor^{LO}$ signature (Pvalue = 0.028, Logrank test).

**Extended Data Fig. 3. Development and validation of the InTumor signature. a.** Proportion of variance for the identified principal components. **b.** Flow-chart of the development of the InTumor signature. **c.** Quantification of CD8+ cells in PDX by IHC in five PDX samples. The negative and positive control samples for comparison are from melanoma cell lines grown in mouse (negative control) or grown in mouse and challenged with Adoptive T-Cell Transfer (ACT) of T-cell receptor-matched cytotoxic T cells[21]). **d.** Examples and quantification of IHC staining of mouse-specific vimentin in PDX samples. The percentage of mouse cells in the PDX are ordered according to the InTumor signature values. Dotted areas indicate mouse cell compartments. Scalebar = 100uM **e.** Examples and quantification of IHC staining of mouse specific PD-L1 in PDX samples. The percentage of mouse cells positive for PD-L1 are ordered according to the InTumor signature values. Dotted areas indicate mouse cell compartments with positive staining for PD-L1. Scalebar = 100uM. **f.** Correlation of the MCPcounter[22] gene sets with the InTumor signature. Metastatic melanoma samples from the TCGA database were used.

**Extended data Fig. 4. PDX-derived cell lines maintain expression of the tumor-intrinsic gene signatures**

**a**. Graphic representation of generation and nomenclature of PDX-derived cell lines. **b.** Heatmap showing the expression of the InTumor genes in PDX samples and matching PDX cell lines.

**Extended data Fig. 5. InTumor signature validation in patient samples. a.** InTumor signature genes plotted according to the ratio Tumor/TME. Top part shows the 14 genes with a 10-fold higher expression in tumor cells over TME genes as measured in scSeq data. **b**. Expression levels of the InTumor signature genes with response to anti-PD-1 in the Riaz et al. dataset. **e.** ROC curves for the InTumor signature, INFg-signature, Cytolytic score and the IMPRES signature in the Riaz et al. patient cohort. The Area Under the Curve (AUC) is indicated behind the signature name. **d.** Response rates to anti-PD-1 separated into $InTumor^{HI}$ and $InTumor^{LO}$ groups (Riaz et al. dataset). e. Tumor mutational burden (TMB) for responders (green) and non-responders (red) split into the $InTumor^{HI}$ and $inTumor^{LO}$ groups.
**f.** Tumor mutation Burden (TMB) compared between $InTumor^{HI}$ and $InTumor^{LO}$ in the Hugo et al. dataset (left panel) and the Riaz et al. data (right). **g.** Relationship between the expression of the InTumor signature genes with response to anti-CTLA-4.

**Extended data Fig. 6. *In vivo* validation InTumor signature. a.** InTumor signature values for a set of 20 common melanoma cell lines. Indicated are the 4 cell lines that were challenged in vivo with ACT. **b.** Growth curves for melanoma cell lines D10, A375, BLM and SkMel-147. Mice were either treated with MART-1 T-cells recognizing tumor (orange) or with Control T-cells (blue). **c.** IHC staining of human specific CD8 for melanoma cell lines D10, A375, BLM and SkMel-147 after challenge with ACT.

**Extended data Fig. 7. InTumor and ExTumor signatures predict response to combination ICB when applied in conjunction. a-b.** Response quadrant and ROC curve and area under the curve (AUC) for the response prediction of the InTumor signature alone. Blue area in the response quadrant indicates samples predicted to respond to anti-PD-1. **c-d.** Response quadrant and ROC curve and area under the curve (AUC) for the response prediction of the ExTumor signature alone. Blue area in the response quadrant indicates samples predicted to respond to anti-CTLA-4.
e. ROC curve and area under the curve (AUC) for the response on the InTumor and ExTumor signatures in conjunction (CA209-038 dataset). **e.** Performance of the IMPRES, IFNg, cytolytic score and the InTumor and ExTumor signatures applied in conjunction on the CA209-038 patient samples. **f.** The InTumor and ExTumor signatures used in conjunction on the samples from the OpACIN clinical trial. Green circles indicate patients who responded to combination ICB therapy, the red circles indicate patients who did not respond to combination ICB. The three blue quadrants indicate InTumor and ExTumor signatures value for samples that are predicted as responders to either anti-CTLA-4, anti-PD-1, or both. The orange quadrant indicates the InTumor and ExTumor signature values for samples that are predicted as non-responders to anti-CTLA-4 as well as anti-PD-1. **g**. Performance of the IMPRES, IFNg, cytolytic score and the InTumor and ExTumor signatures applied in conjunction on the OpACIN patient samples. **h.** Difference in tumor mutational burden (TMB) for predicted responders and predicted non-responders to combination therapy in the OpACIN dataset. **i.** Kaplan-Meier plot comparing recurrence-free survival (PFS) for patients predicted as responders (blue) to patients predicted as non-responders (orange) for the OpACIN trial.

**Extended data Fig. 8. Comparison of groups from the response quadrant.** Percentage of patients with a clinical response in the four groups in the response quadrant separated into three groups (anti-CTLA-4, anti-PD-1 and combination of anti-CTLA-4 + anti-PD-1 treated).

**Extended data Fig. 9. SNP comparison within the set of PDX samples.** Heatmap representing the percentage overlap in SNPs between PDX samples (n=95, left bottom), between PDX cell lines (n=22, right top) and between PDX samples and PDX cell lines (left top). Samples with the same parental origin are colored.

#### Extended Data Fig. 1

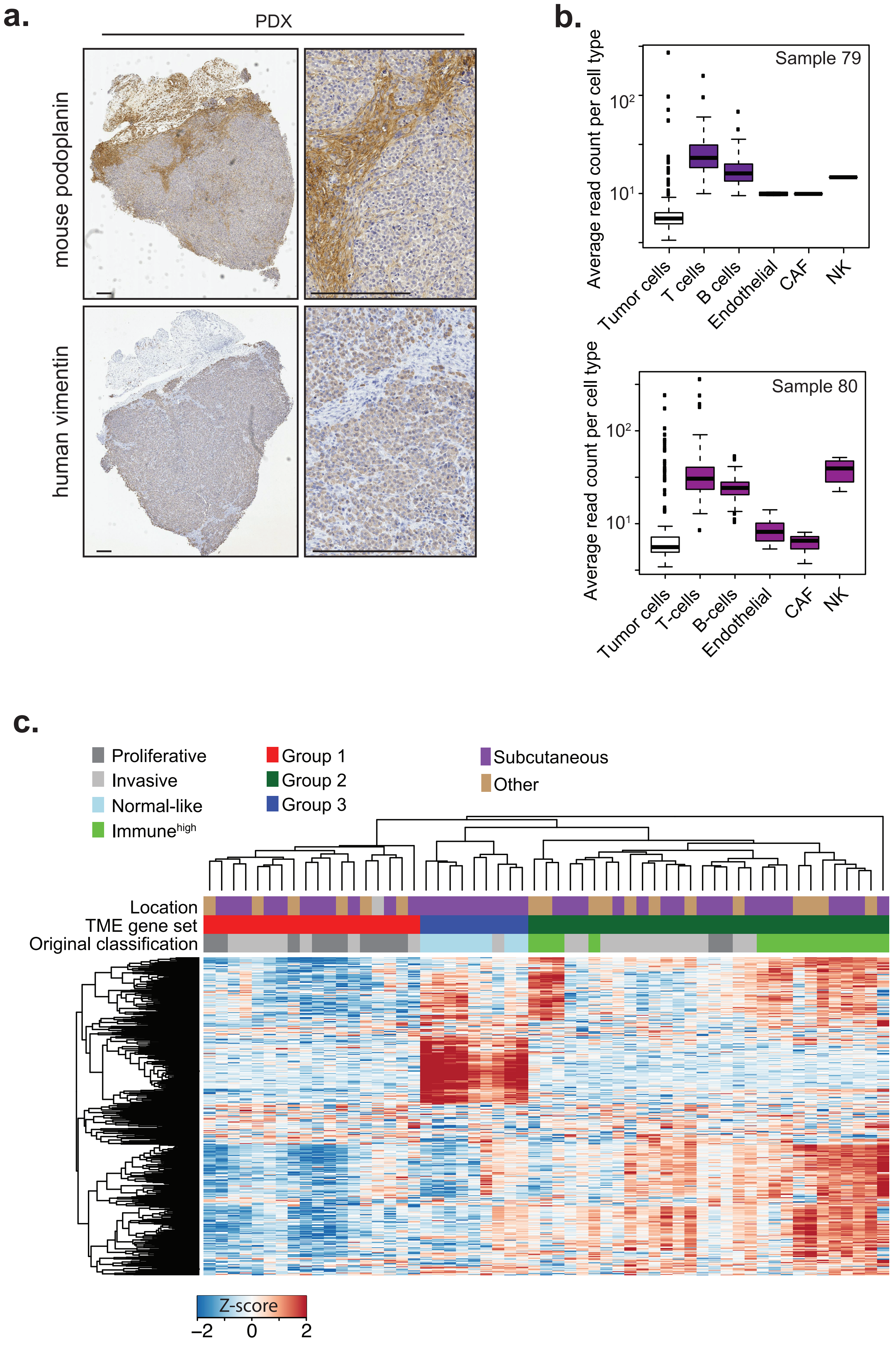

### Extended Data Fig. 2

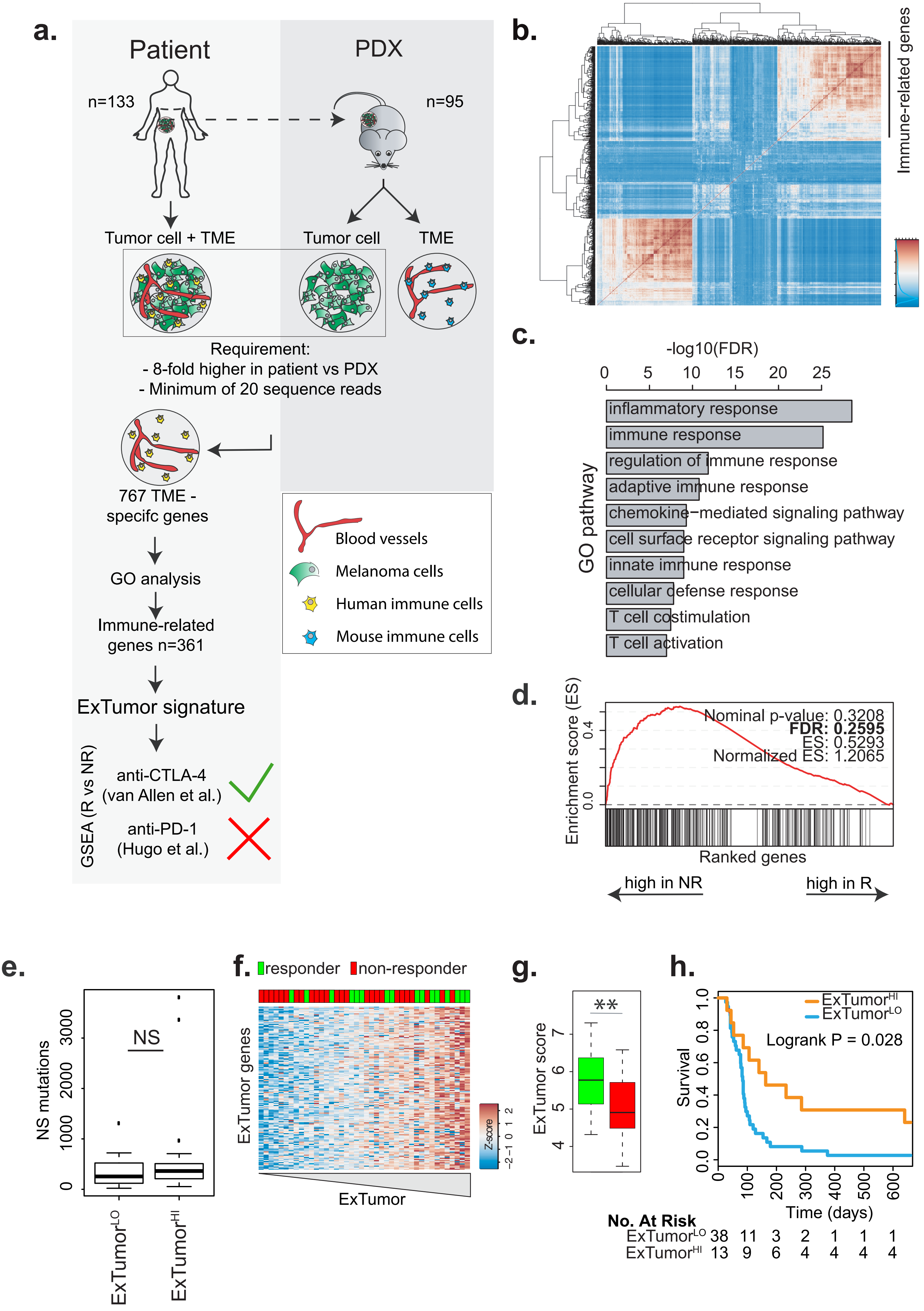

### Extended Data Fig. 3

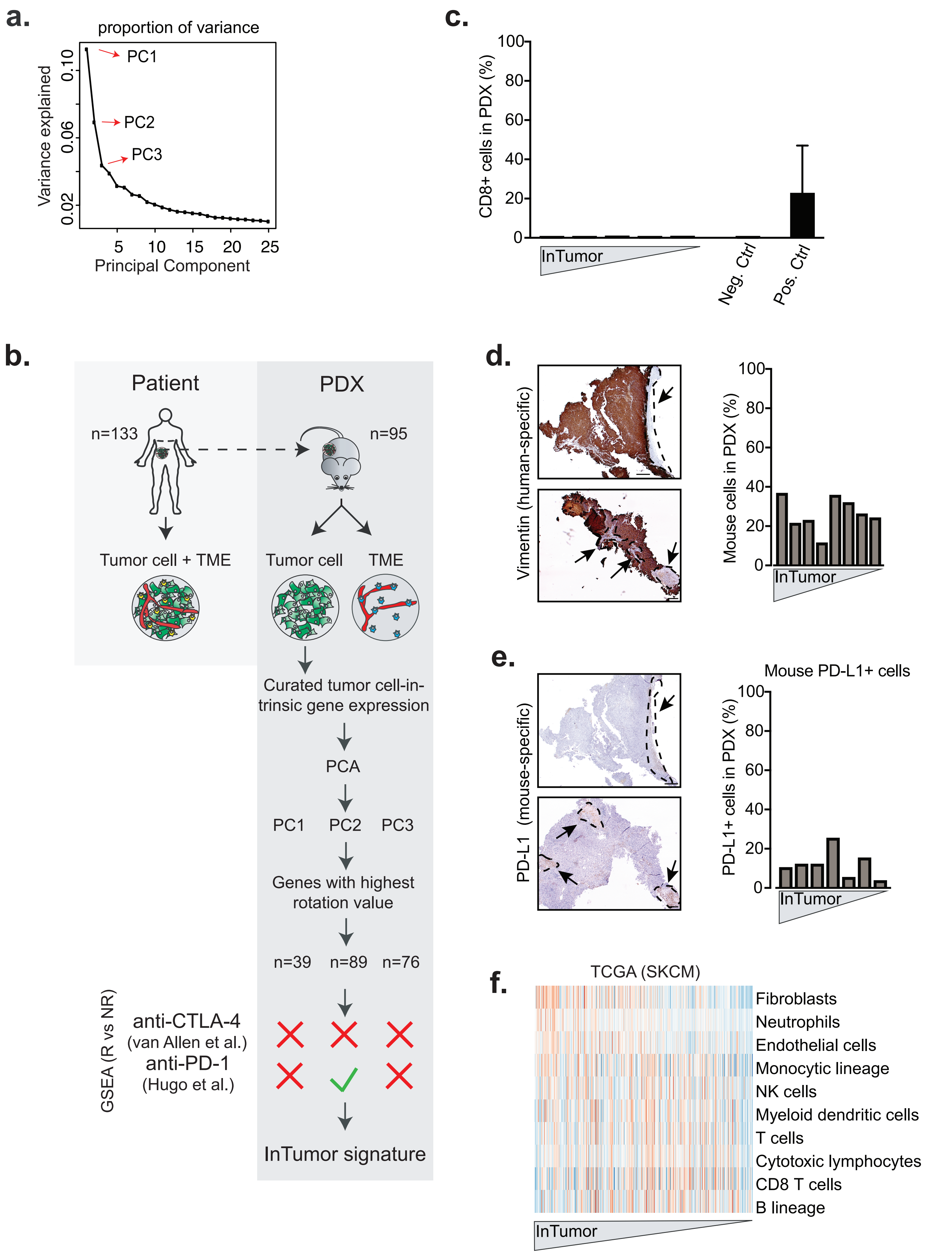

### Extended Data Fig. 4

**a.**

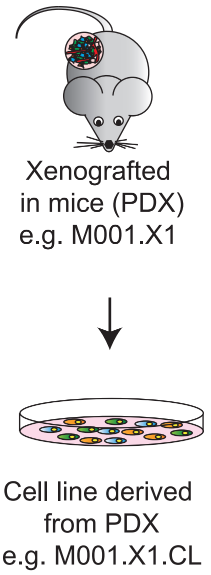

**b.**

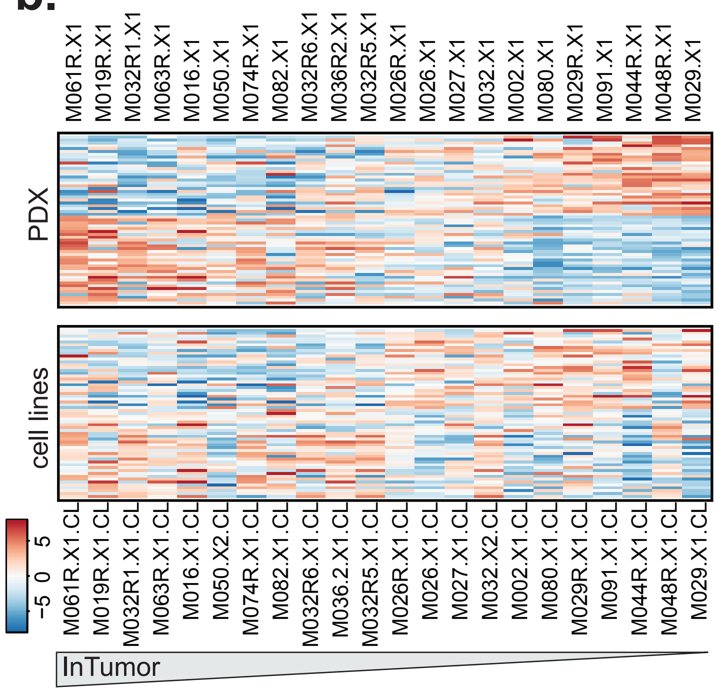

### Extended Data Fig. 5

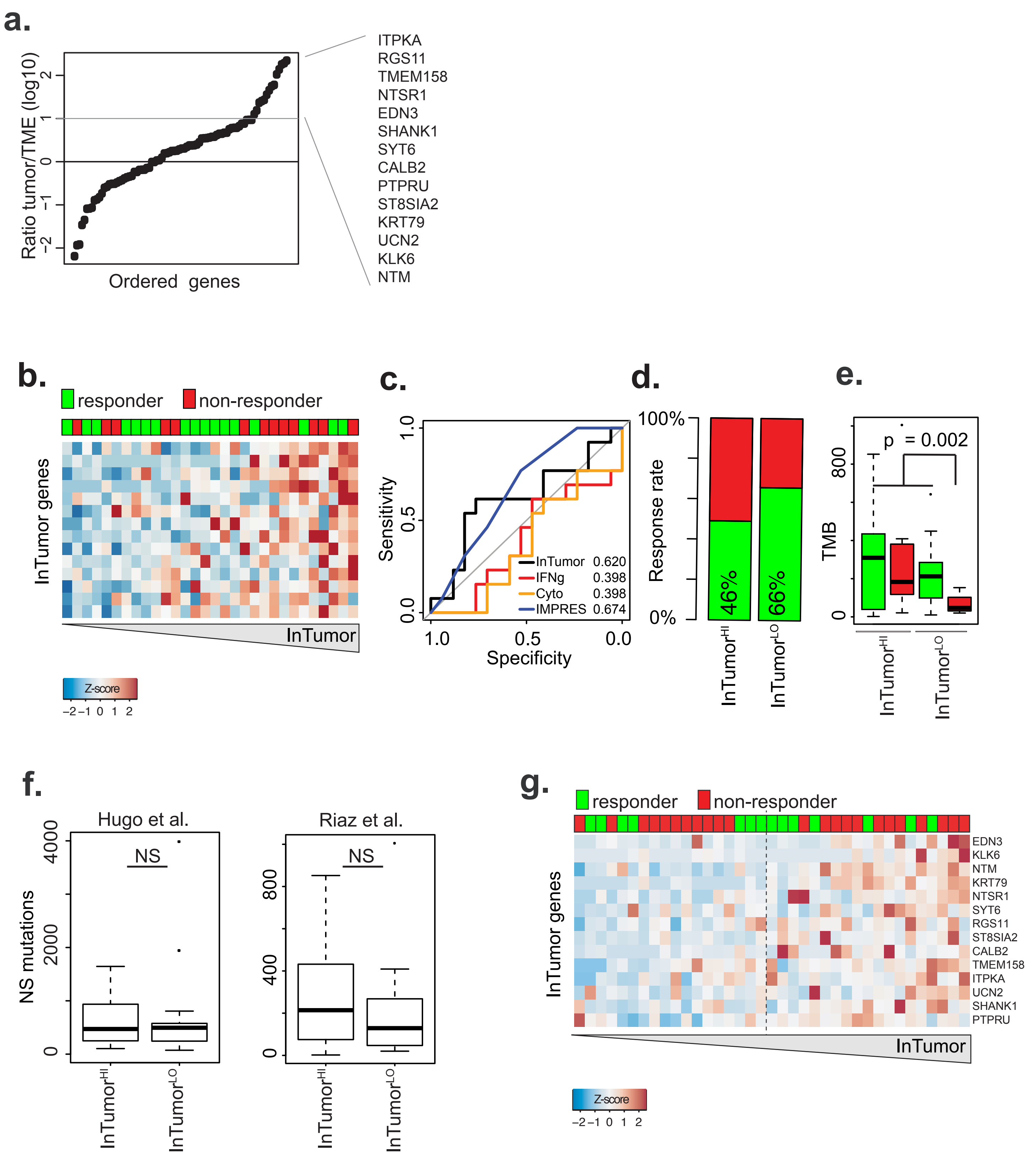

### Extended Data Fig. 6

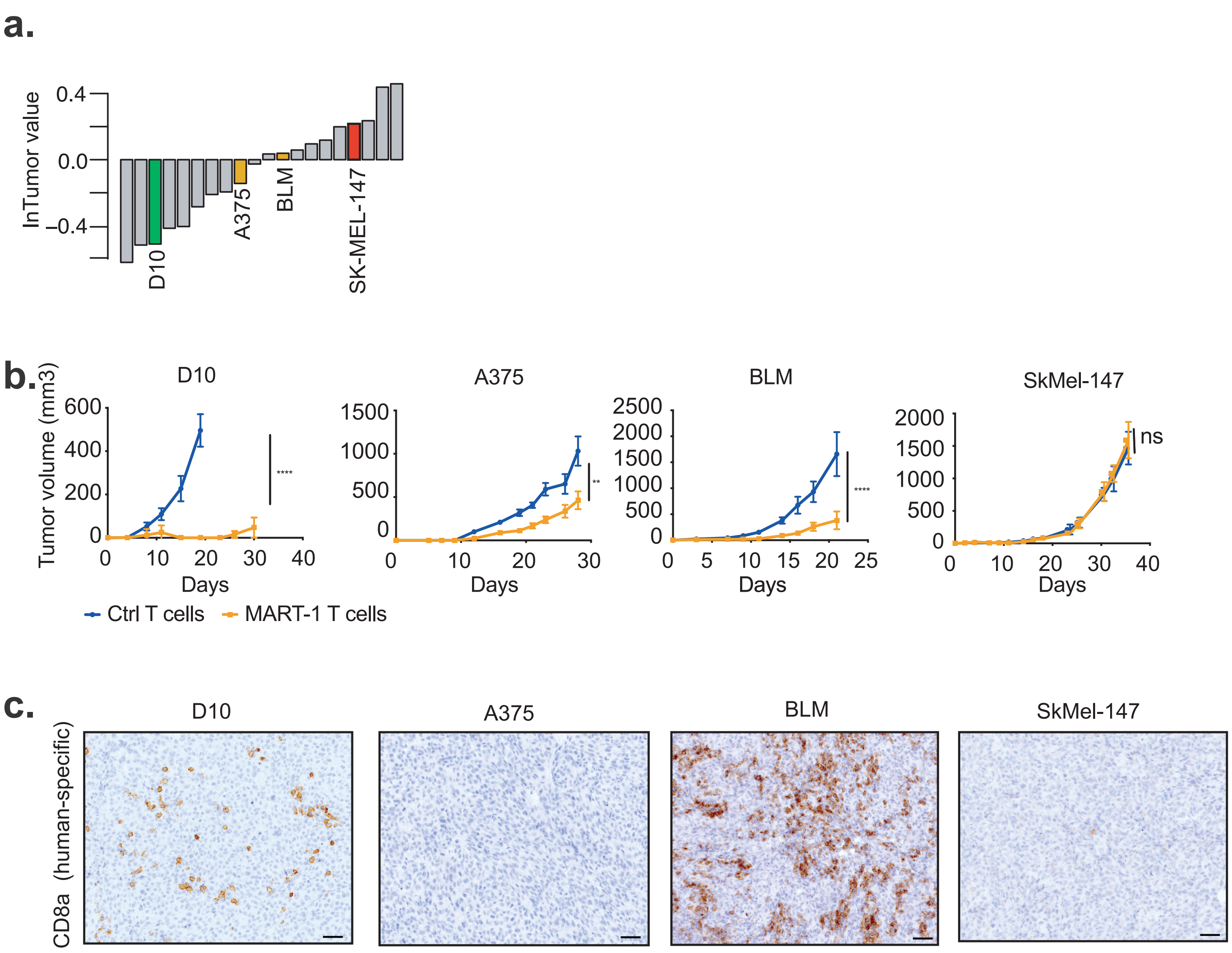

### Extended Data Fig. 7

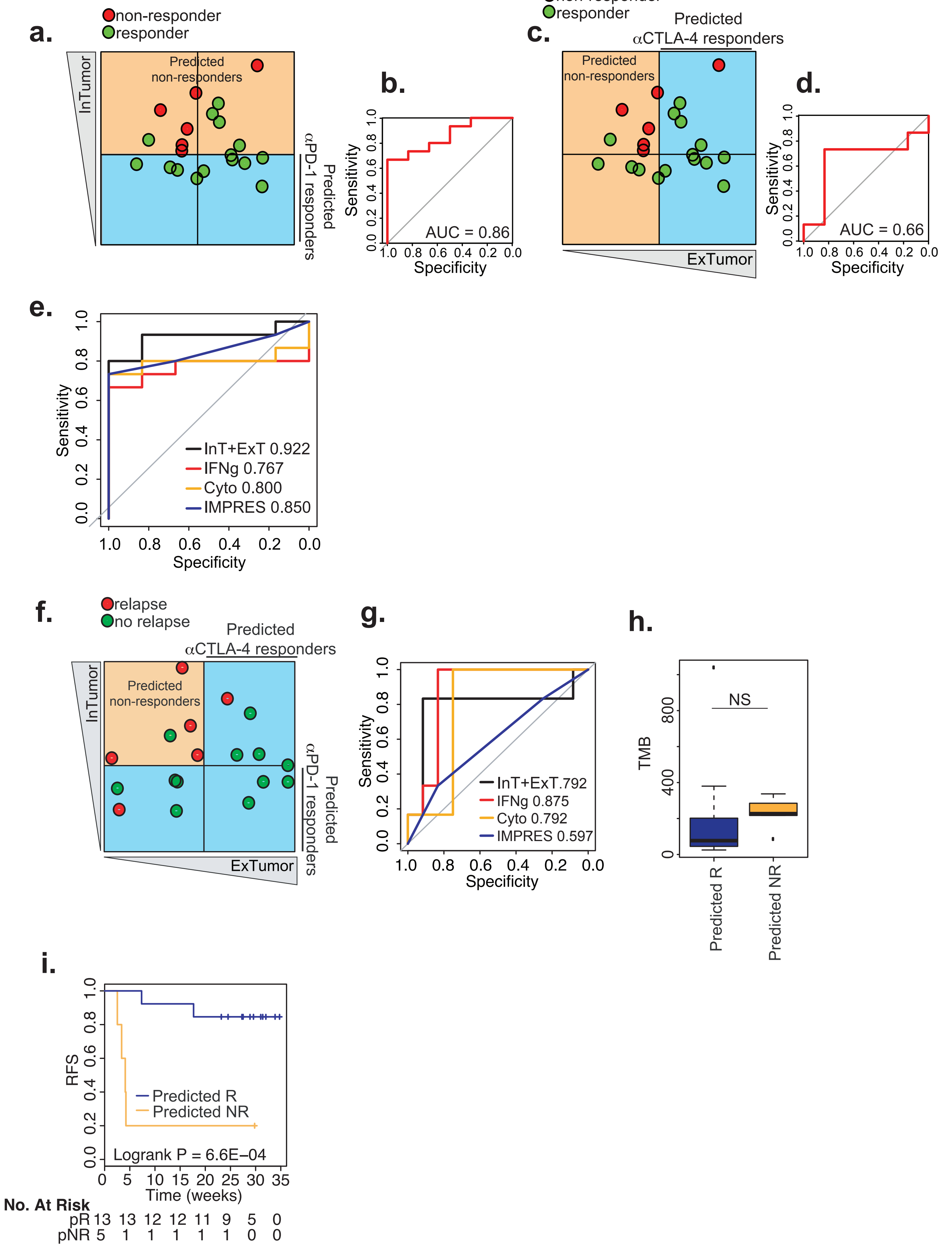

### Extended Data Fig. 8

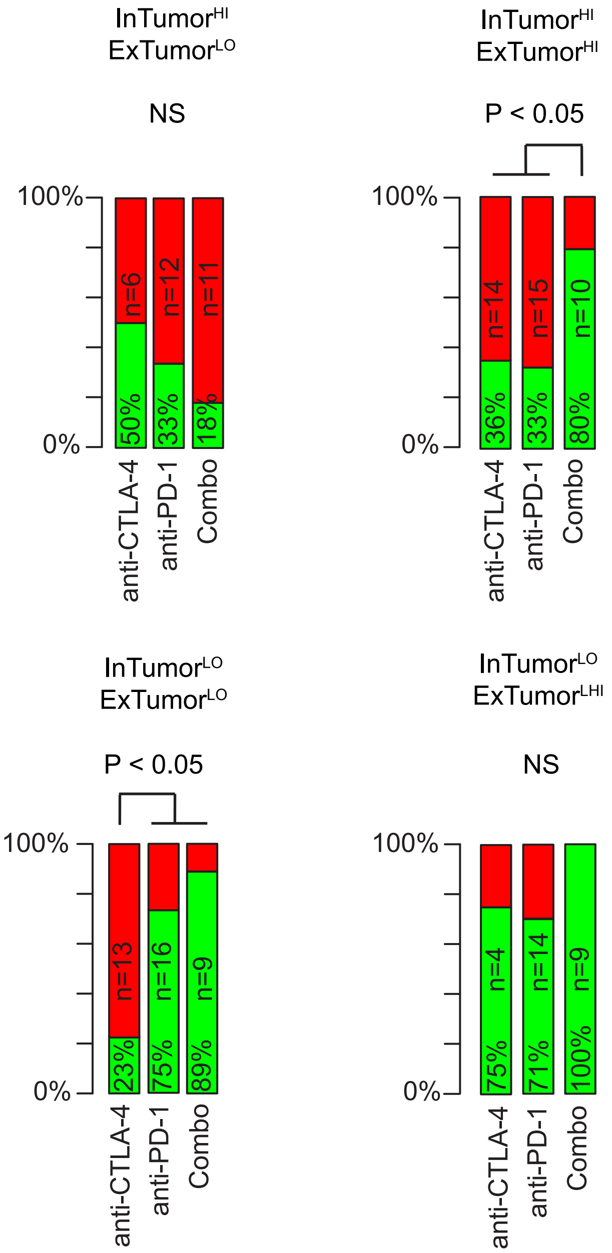

### Extended Data Fig. 9

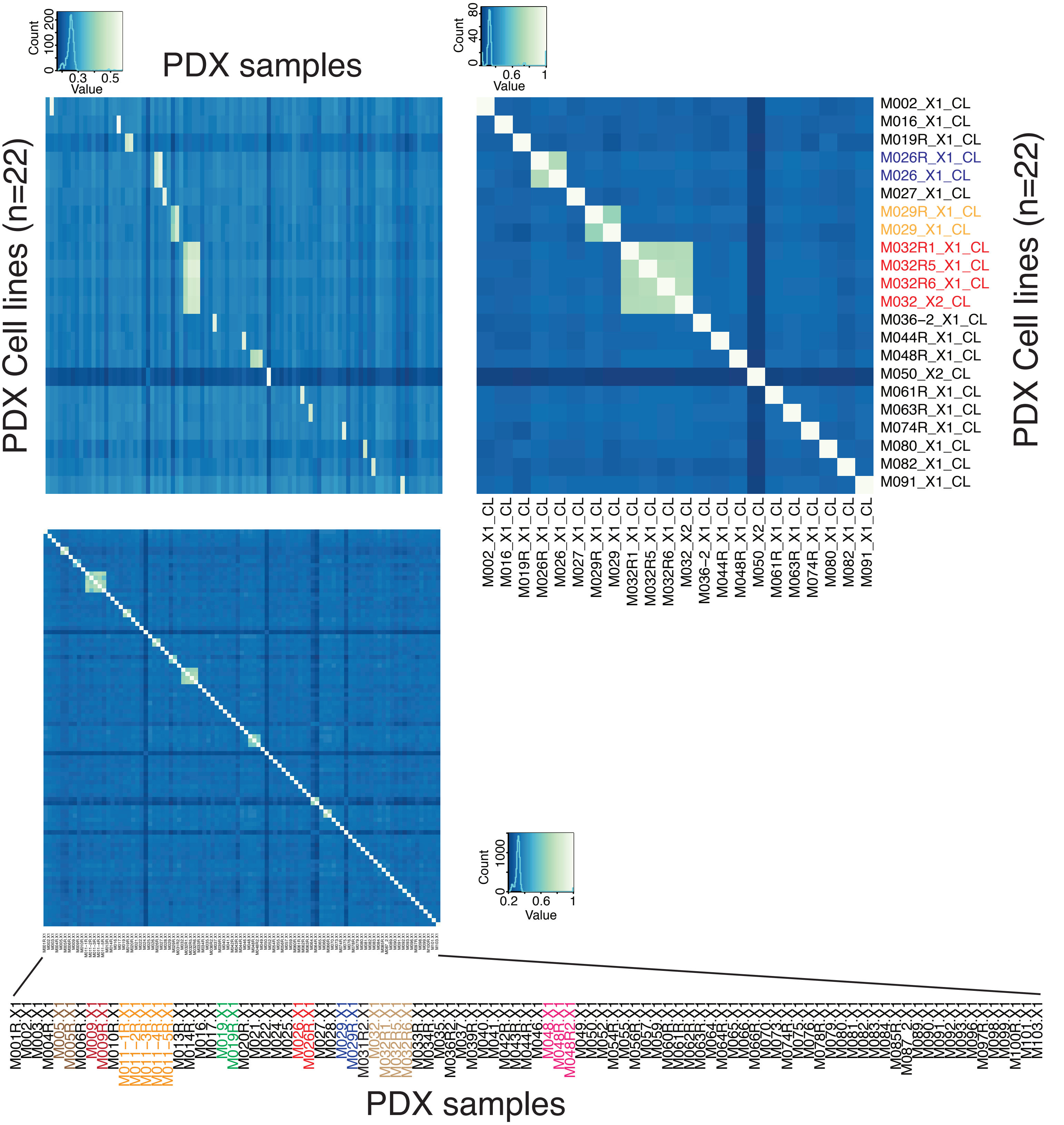
